# An Order of Magnitude Signal-to-Noise Improvement of Magnetic Resonance Spectra using a Segmented-Overlap Fourier-Filtering and Averaging (SOFFA) Approach

**DOI:** 10.1101/2025.04.21.649794

**Authors:** Jason W. Sidabras

## Abstract

Segmented-Overlap Fourier-Filtering and Averaging (SOFFA) data acquisition method is described in detail for magnetic resonance spectroscopy. In this work the four processes that encompass the SOFFA data acquisition method are detailed: (i) oversampling spectral segments, (ii) Fourier block-filtering, (iii) segment-overlap averaging, and (iv) decimation. Three experimental examples are shown. Conventional Continuous Wave (CW) Electron Paramagnetic Resonance (EPR) is compared to SOFFA-CW of a single reduced [4Fe-4S]^+^ (S=1/2) at concentrations of 1 mM, 100 *µ*M, and 10 *µ*M showing an average increase in concentration sensitivity by a factor of 5.6. Experimental comparison of CW and SOFFA nonadiabatic rapid scan (SOFFA-NARS) data with similar filter parameters and field-modulation amplitude demonstrates a factor of 10.3 in signal-to-noise improvement for a 150 *µ*M sitedirected spin-labeled Hemoglobin in 82% glycerol at 18°C. Conventional and SOFFA rapid scan (SOFFA-RS) data was collected for a minuscule amount of lithium phthalocyanine (LiPC) and a factor of 9.6 in signal-to-noise improvement is demonstrated. The signal-to-noise improvements were made for the same data acquisition times on standard commercial instruments. This method can be implemented to perform real-time segmented processing and, combined with more sophisticated averaging methods, will push the state-of-the-art sensitivity in magnetic resonance spectroscopy.

## 1. Introduction

In this work, the Segmented-Overlap Fourier Filtering and Averaging (SOFFA) method for magnetic resonance data acquisition and filtering is established. In a typical continuous-wave (CW) Electron Paramagnetic Resonance (EPR) experiment a spectrum is collected as a continuous signal, *i.e*. the spectroscopist must “play” it until the end and, only then, repeat for averaging. However, the signal is typically time invariant, and therefore can be collected in a number of unique ways. It is shown herein that by collecting a magnetic resonance spectrum in segments an additional parameter related to the overlap of each scan is introduced. Each acquired segment contains field- or frequency-stepped correlated signal and uncorrelated noise. For processing, each segment is oversampled, filtered block-wise, and concatenated resulting in a signal-to-noise improvement compared to traditional CW and rapid scan data collection, averaging, and filtering methods for the same measurement time. The SOFFA data acquisition method can be adopted for many magnetic resonance experiments and is demonstrated for conventional CW and rapid scan techniques. The scheme is illustrated in Fig. 1.

**Figure 1:**
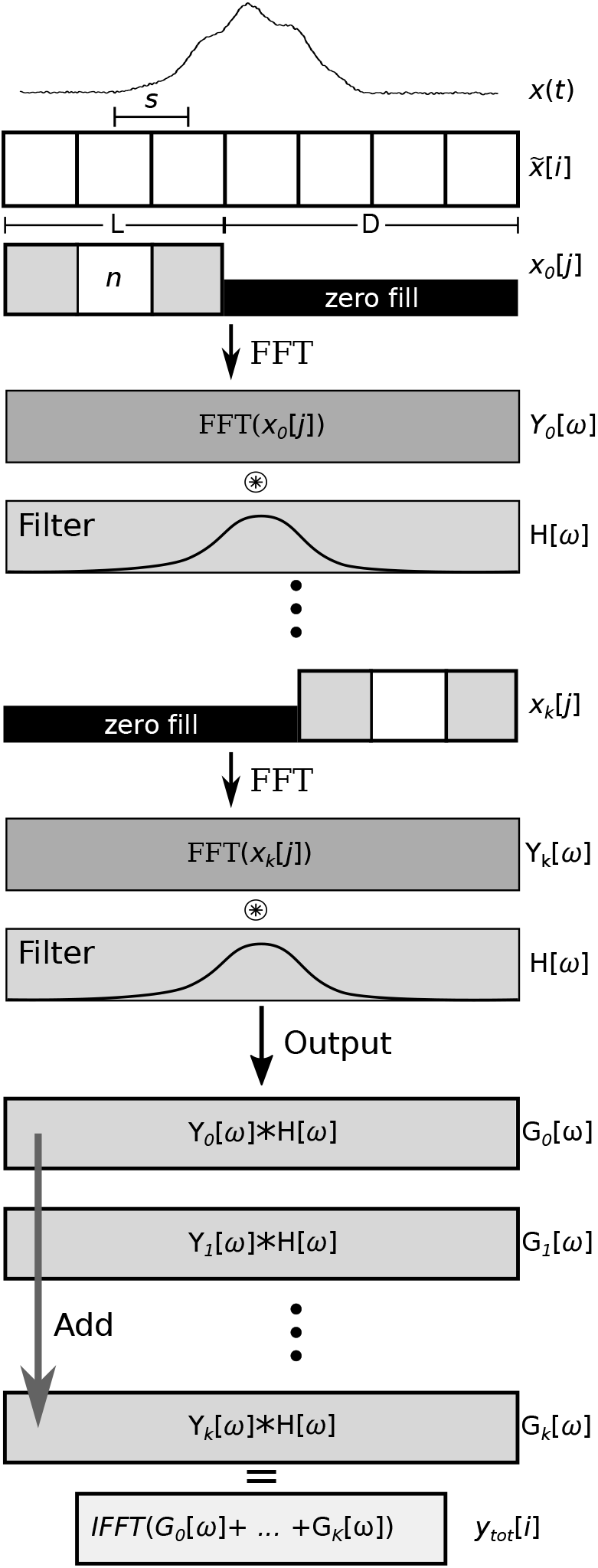
Illustration of the SOFFA data acquisition method used to concatenate and filter segmented magnetic resonance data. Herein, the filter H[*ω*] is a Gaussian in the frequency domain. The collected segments *x*_*k*_[*j*] and final filtered function *y*_tot_[*i*] are related to the segmented sweep width in mT or MHz, for field or frequency sweep, respectively.

The improved signal-to-noise ratio (SNR) achieved by the SOFFA method can also be leveraged to reduce the time required for an experiment, thereby increasing throughput. In traditional EPR experiments, the SNR is often improved by averaging multiple scans, which can be time-consuming. However, with the SOFFA method, the same level of SNR can be achieved in a shorter amount of time, typically by the square of the number of traditional averages. This means that the experiment time can be reduced, allowing for more samples to be analyzed in a given time period. By using the SOFFA method, this relationship can be decoupled, allowing for improved SNR without a corresponding increase in experiment time. This can have significant implications for high-throughput applications, such as screening large numbers of samples or performing kinetic studies.

The non-adiabatic rapid scan (NARS) [1, 2, 3, 4, 5] and adiabatic rapid scan (RS) [6, 7, 8, 9, 10, 11] methods collect pure-absorption EPR spectra using fast field or frequency sweeps and modern digital signal processing. Both rapid scan methods collect quadrature pure-absorption EPR spectra, which can be pseudo-modulated to the conventional first-derivative EPR spectrum.

In a RS experiment, the field or frequency is swept at a rate and amplitude that is larger than the relaxation rate of the spin system. The RS experiment bridges the gap between CW and pulse EPR experiments and is categorized as a passage experiment. The RS EPR signal is similar to a Free Induction Decay (FID) signal and must be Fourier transformed and deconvolved to regain the slow-scan EPR pure-absorption signal. [6] The advantage of a RS experiment is the field or frequency can be swept at such rate and amplitude that the spin system power saturation characteristics are shifted to higher power. Therefore, the input power can be increased to keep ∂*B*_1_*/*∂*t* constant. Improvement of the signal-to-noise by over an order of magnitude is feasible. RS has been adopted to many experiments, such as spin-labeled nitroxides [8], *γ*-irradiated materials [12, 13], EPR imaging [10], and quantitative EPR. [11] Typically, RS experiments employ large amplitude (upwards of 20 mT) and rates (MHz or higher) for field or frequency sweeps.

In a NARS experiment, the field or frequency is swept at a rate and amplitude that allows the spin system to return to thermal equilibrium and no passage is observed. In this sense, NARS mimics a CW EPR experiment and the power saturation profile of the sample remains unperturbed. Typically, NARS experiments employ low amplitude (1 mT) and rate (below 50 kHz) for field or frequency sweeps. In order to collect a full EPR spectrum, the static field is stepped and data is collected. Each step is windowed, zero filled, concatenated, and filtered. The advantage of NARS comes from the modern data collection methods (averaging, digital filtering, etc.), collection of pure-absorption spectra, and signal-to-noise improvements due to overlapping segments during concatenation. NARS has been used by Hyde and colleagues [1, 3] at L-band (1.2 GHz) to extend the deconvolution method of CW EPR distance determination from 18-25 Å to 30-35 Å [2] and increase the resolution for a copper imidazole complex. [4] Additionally, NARS was independently shown by Eaton and colleagues in applications of spectra as wide as 620 mT. [5]

It has been established that non-adiabatic and adiabatic rapid-scan data collection produces a comb filter that suppresses high frequency noise. [14, 15] However, these methods are susceptible to low frequency noise that is correlated with the field sweep or the transfer function of the frequency response of the system. Filtering such noise is problematic since the features may contain the same frequency content as the absorption spectrum. Methods have been proposed for the background correction of RS data [7] and can be easily combined with the deconvolution needed for sinusoidal sweeps. [16, 17] Additionally, pseudo-modulation using the Moving Difference (MDIFF) algorithm to calculate the conventional first-harmonic derivative-like spectrum further reduces some low frequency noise. [4] However, the challenge of properly filtering NARS and RS data still remains. The SOFFA data acquisition method is shown to remove challenging noise profiles and that the method is well suited for magnetic resonance spectroscopy with minimum changes to data acquisition.

The SOFFA data acquisition method is tested with (i) an experimental comparison between an averaged and filtered CW spectrum and a SOFFA-CW spectrum using concentrations of 1 mM, 100 *µ*M, and 10 *µ*M of *apo* [FeFe]-hydrogenase (single reduced [4Fe-4S]^+^; S=1/2; Ref. [18]), (ii) an experimental comparison of a SOFFA-NARS spectrum of 150 *µ*M site-directed spin-labeled Hemoglobin protein in 82% glycerol at 18°C to averaged and filtered CW spectrum, and (iii) an experimental comparison between traditional and SOFFA-RS spectra of a minuscule amount of lithium phthalocyanine (LiPC). For direct comparison the measurement time was kept constant for each set of experiments.

## 2. Methods

All data is processed using Wolfram Mathematica (v. 11.3; Champaign, IL, USA) where the SOFFA algorithm was implemented and tested for a number of experiments. Data was collected on various spectrometers with parameters as specified. To collect a segmented-CW EPR spectrum a ProDEL program was written for Bruker Xepr where the field was stepped and the data was collected.^1^ Each step was placed in a 2D representation and saved in typical Bruker format (DSC/DTA).

Here the nomenclature of the Welch FFT procedure for power spectral density is adopted, due to the similarity of segmentation of a discretized signal, with variables illustrated in Fig. B1. [19] The data is processed in the following manner, as illustrated in Fig. 1:

1. Data is collected at a series of field steps *s* in mT.
2. The data is discretized and windowed to the appropriate length *L*. The length *L* is determined by the number of points in a discretized field-step *n* (shift-step size) multiplied by the overlap factor *m*. The discretized shift-step size *n* can be estimated by

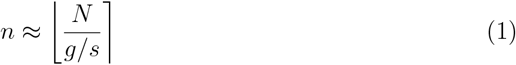

where *N* is the number of points in the sweep (e.g. 4096), *g* is the field sweep length in mT, and *s* is the field step in mT. The length of the window *L* and the number of points in a sweep *N* can be equal, but the windowing can be independent of the total number of points if edge effects (i.e. rapid scan turning points) need to be eliminated. The ratio of the field sweep length and step size yields a maximum overlap value *m*. The shift-step size *n* is rounded to the nearest integer. The integer *m* is a multiplier of *n* used for segmental overlap, generating *x*_*k*_[*j*], where *x*_*k*_ is a dataset with point *j* = 0, 1, 2, …, *L* − 1 and *k* is the index of total number of segments *K*. The number of segments are calculated by

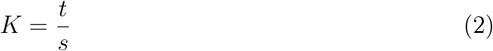

where *t* is the total spectral sweep width in mT. The ratio is chosen such that K is an integer. No overlap exists at *m* = 1 and symmetric overlap requires *m* as an odd number.
3. The data is zero filled with *D* points to the total discretized length of 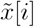 with *D* being step-shifted by *n* to align the segments by absolute field.
4. The data is Fourier transformed (FFT) and, in this work, a fixed width Gaussian filter is applied, defined by

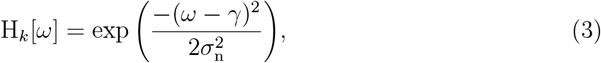

where *ω* is the discretized frequency, *γ* in the central point in the Fourier transformed data and has a variance of 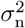.The smallest variance 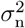 which shows no spectral broadening is chosen for each *L*.
5. Filtered data is concatenated to yield *G*_tot_[*ω*] which is Inverse Fourier transformed (IFFT) resulting in a filtered whole spectrum *y*_tot_[*i*]. The spectrum is truncated to remove the head and tail (data without overlap) by removing the first and last *L* points and *y*_tot_[*i*] is decimated to a conventional number of points, such as 4096.

Signal-to-noise is calculated by the ratio of the acquired signal mean to the standard deviation of the noise voltage, defined by

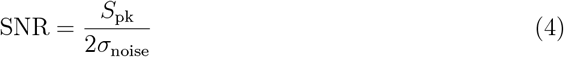

where *S*_pk_*/*2 is the mean value of the peak-to-peak (or peak for absorption spectra) signal, and *σ*_noise_ is the standard deviation of the noise. [20, 21] The standard deviation of the noise is calculated in Mathematica using points in an off-resonance region.

### 2.1. Filtering Tests of Simulated Data

Simulated data of a Gaussian was used to represent an absorption spectrum. Two simulations were performed to test the SOFFA algorithm. The first simulation used strictly random (white) noise and was generated in Mathematica using the `RandReal` function with a *max* and *min* range defined in such a way that the signal-to-noise ratio was approximately 4 for a single unfiltered scan. The `RandReal` function is a uniformly distributed pseudo-random number generator, see Fig. A1(*vi*). The second simulation used frequency-dependent (pink) noise created in Mathematica using the `AudioGenerator[“Pink”]` function to generate pseudorandom frequency-dependent noise. This noise was added to a white noise which established a noise floor, see Fig. A1(*v*), such that the signal-to-noise of a single unfiltered scan was 10. Pink noise was then added to decrease the signal-to-noise to approximately 4 for a single unfiltered scan.

Each data point in the simulation was repeated 100 times to obtain statistically converged results. For simulated spectra, the 95% confidence interval is shown as an indication of the precision of the estimated mean. Each point was tabulated and plotted alongside conventional averaging for comparison. Segmented data was generated by stepping a window across a noiseless spectrum and generating noise at each step. Each step had an *n* = 20 and 6000 points. A total of 500 steps were generated over the spectrum. The overlap *m* was varied and each point was either filtered after concatenation and decimation (Post-Add Filter; Methods of Ref. [4]) or filtered in Fourier space and concatenated (SOFFA). For the continuous-wave simulation the unfiltered (Raw) data had 4096 points and a low effective time-constant. The data was then filtered using the same Fourier transform Gaussian filter. This was repeated for both white and pink noise.

The filter variance 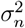 was chosen as small as possible without showing spectral broadening. For conventional averaging, the spectral bandwidth of the signal remained constant and a standard deviation of *σ*_*n*_ = 35 was used. Similarly, the spectral bandwidth for filtering after decimation (Post-Add Filter) is constant and a standard deviation of *σ*_*n*_ = 30 was used. For the SOFFA method, the spectral bandwidth is a function of *L* and, therefore, must be found for each overlap. Values are tabulated in Table 1.

**Table 1:** Comparison of signal-to-noise values using white (SNR_W_) and pink (SNR_P_) noise with traditional averaging and filtering (Trad. Ave.), Post-Add Filter processed using the methods of Ref. [4], and the methods introduced here (SOFFA) for a Gaussian absorption spectrum.

| $m_a$ | Trad. Ave. | | | $m$ | Post-Add Filter | | | SOFFA | | |
| --- | --- | --- | --- | --- | --- | --- | --- | --- | --- | --- |
| | $\sigma_n$ | $\text{SNR}_W$ | $\text{SNR}_P$ | | $\sigma_n$ | $\text{SNR}_W$ | $\text{SNR}_P$ | $\sigma_n$ | $\text{SNR}_W$ | $\text{SNR}_P$ |
| 1 | 35.0 | 33.0 | 9.1 | 1 | 30.0 | 32.9 | 14.78 | 25.0 | 71.5 | 15.2 |
| 5 | — | 73.9 | 21.2 | 5 | — | 80.2 | 18.1 | 20.0 | 192.6 | 22.1 |
| 10 | — | 104.5 | 29.1 | 10 | — | 108.4 | 22.5 | 17.5 | 292.4 | 27.2 |
| 20 | — | 147.8 | 39.1 | 20 | — | 155.1 | 32.0 | 20.0 | 386.6 | 35.5 |
| 30 | — | 181.0 | 48.7 | 30 | — | 184.9 | 41.1 | 20.0 | 520.7 | 45.8 |
| 50 | — | 233.6 | 64.1 | 50 | — | 246.4 | 55.9 | 22.5 | 625.7 | 62.6 |
| 100 | — | 330.4 | 92.2 | 100 | — | 349.4 | 84.7 | 27.5 | 963 | 97.2 |
| 200 | — | 467.3 | 131.1 | 200 | — | 491.8 | 128.0 | 45 | 1101 | 152.3 |
| 300 | — | 573.1 | 157.7 | 300 | — | 675.2 | 164.6 | 70 | 1092 | 213.0 |

## 3. Theory

In this work, linear noise is assumed, such that,

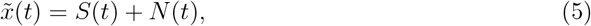

where *S*(*t*) is the signal that is detected and *N* (*t*) is either random (white) or frequencydependent (pink) noise. Noise that is a function of the signal is beyond the scope of this work.

For conventionally acquired data, the number of points, scan rate, and time constant are chosen to represent the spectrum completely and without spectral distortions, simulated data for visualization is shown in Fig. 3A. For each new scan, the noise is uncorrelated, therefore, the data can be averaged and filtered. For example, filtered data of Fig. 3A is shown in Fig. 3C. Fourier filtering averaged data before or after summing the individual scans is equivalent. With traditional averaging, the signal-to-noise follows a strict 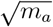 gain,

**Figure 2:**
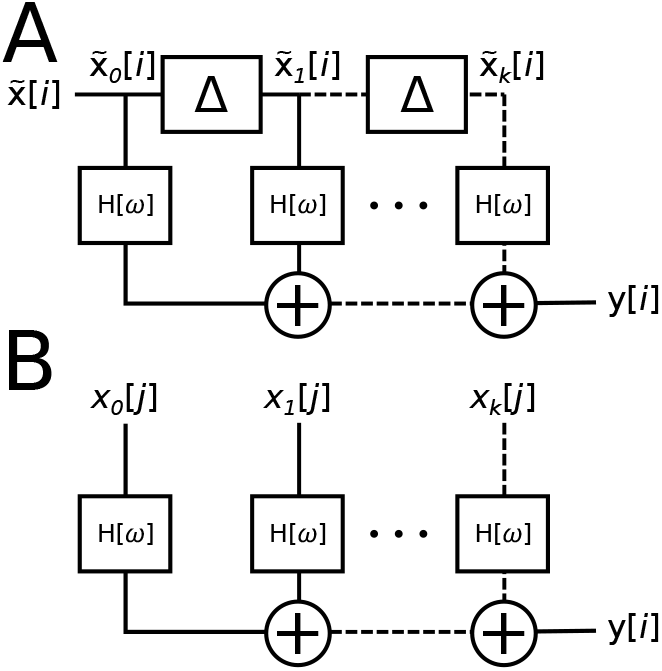
(A) The Overlap-Add filter design is used for a large time-domain signal *x*(*t*) where it is discretized 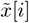and segmented by delayed blocks Δ. Each discretized window is Fourier filtered by H[*ω*] and concatenated. (B) In a segmented magnetic resonance experiment, each overlapped segment has noise that is uncorrelated. Each segment is discretized separately, Fourier filtered by H[*ω*], and overlapping segments are further averaged during concatenation. Without overlap (*L* = 1*n*) the methods of **A** and **B** are equivalent.

**Figure 3:**
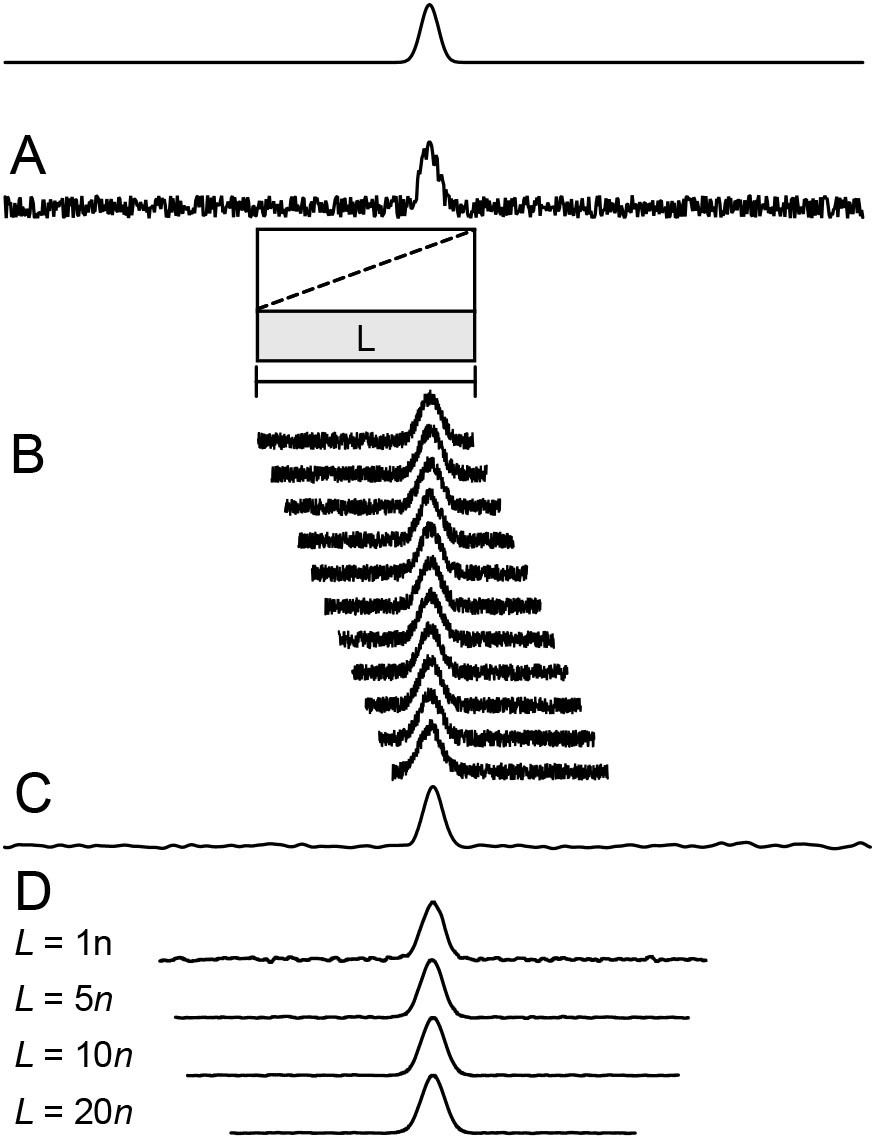
A Gaussian absorption spectrum is simulated with a half width of 1 mT and an amplitude of unity. Shown here is a representation of the simulated experiment. (A) Random (white) noise is added. (B) Small set of unfiltered segmented data is plotted showing field steps *s*. Illustration of segmentation is shown in the inset. (C) When a Gaussian convolution filter of Eq. 3 is applied to **A**, the expected improvement in signal-to-noise is shown, similar to increasing the time constant. (D) However, as *L* is increased, further signal-to-noise improvement is exhibited due to the concatenation of filtered data per the SOFFA method. This experiment is repeated 100 times and the mean of the signal-to-noise ratios are plotted for white and pink noise, Figs. 4A and 4B, respectively. white and pink noise data is tabulated in Table 1.

where *m*_*a*_ is the number of averages. The same signal/time trade-off exists when increasing the number of collected data points while maintaining equivalent measurement time.

If a signal input is infinitely long or for signals that need to be filtered in real time, the Overlap-Add method is used with short-time Fourier transform (STFT) block-filtering for efficient digital signal processing. [21, 22] Block-filtering breaks a long or continuous signal *x*(*t*) into a series of spectral windows with a fixed discrete length. Each window is filtered individually in Fourier space and concatenated back together, as illustrated in Fig. 2A. Finite-impulse response (FIR) filters with block-filtering are easily implemented and will be the focus of this work. Finally, the output of block-filtering and recombining the data with an Overlap-add or Overlap-save method is equivalent to filtering the long signal *x*(*t*) data directly.

In this section the four processes utilized to create the SOFFA data acquisition method are detailed: (i) Oversampling Spectral Segments, (ii) Fourier Block-Filtering each segment separately, (iii) Segment-overlap averaging, and (iv) Decimation. It is the combination of the these processes that improves the signal-to-noise ratio for the same amount of data collection time.

### 3.1 Oversampling Spectral Segments

When implementing a block-filter approach, the discretized continuous input signal must be down-sampled by the number of blocks and then, after filtering, up-sampled and concatenated. [21] The down- and up-sampling ratios do not have to be equivalent and each block could have overlapping segments. [23] For example, if an input signal has 4096 points, a series of 32 blocks can be formed without overlap with *L* = 128. The advantage of employing block-filtering is the ability to process the blocks in parallel, illustrated in Fig. 2A, and speed up the convolution process for real-time processing or for continuous discrete-signals.

When implementing traditional block-filtering, segments are generated by windowing the whole signal where the windowing function with window size *L*, which yields a long signal segmented into *k* segments with the length of *L* (*i* = 0, 1, …, *L*− 1) separated by *n*, which is the discretized window step delay size. Once windowed and Fourier transformed, each discretized signal is considered a Short-time Fourier transform (STFT) and can be filtered in Fourier space with convolution techniques. Since each segment is generated from a single input the noise is correlated and an appropriate windowing function *w*[*i*− *n*] is required to minimize processing noise and aliasing artifacts. [24] When each block is recombined using the Overlap-add or Overlap-save algorithm, the signal-to-noise of the filtered signal is exactly equivalent as if the whole signal was processed directly.

In the SOFFA data acquisition method, each field/frequency swept segment has the following properties: (i) the overlapping noise is non-coherent, (ii) field- or frequency-dependent features are considered signal, and (iii) each segment is an oversampled section of the whole spectrum. In the case of the SOFFA method, all segments are collected separately and can then be further windowed to the desired length to remove potential edge effects the windowing function with window size *L*. Stepping *s* over the spectrum yields *k* individually collected segments. Each segment is offset by the discrete shift-step size *n*. Since magnetic resonance produces a stationary and time-invariant signal, the corresponding output blocks can be zero filled by a length of the total collection size to begin at the same absolute step size, aligning the field or frequency dependent features. The zero-fill parameter is then shift-stepped by *n* to align field dependent features. To maximize the overlap averaging after filtering, the windowing function is chosen to be rectangular and no aliasing issues exist if oversampling is used. Unlike windowing in typical block-filtering, which benefits greatly from window choice [24], a rectangular window will maximize the individual contribution of each segment during the concatenation process. Any higher order frequencies generated by discontinuities are removed in the filtering process and result in proper concatenation and aliasing.

In the SOFFA data acquisition method, each segment is oversampled in order to reduce the noise spectral density while keeping the signal spectral density constant. [21] One can estimate an effective time constant,

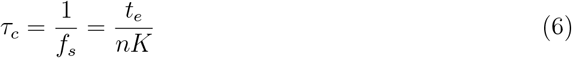

where *f*_*s*_ is the effective sampling frequency, *t*_*e*_ is the total experiment scan time, *n* is the shift-step size and *K* is the total number of segments. If the noise is predominantly white and a perfect low-pass filter is used, the signal-to-noise is improved by the oversampling ratio

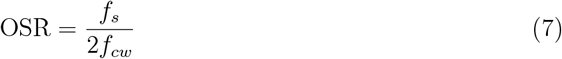

where *f*_*cw*_ is the inverse of the time constant *τ*_*cw*_ of the conventional CW data collection.

Signal-to-noise is improved in oversampling by reducing the spectral bandwidth by the OSR and effectively spreading noise power spectral density by the same amount, see Fig. A1 (*ii* and *iii*). [21] Since segmented magnetic resonance spectra are collected, a further decrease the spectral bandwidth is exhibited since only a fragment of the spectrum is sampled.

Once each of the segments is sufficiently oversampled, noise-shaping can be employed to reduce the high frequency content outside the spectral bandwidth by block-filtering techniques.

### 3.2 Fourier Block-Filtering each Segment Separately

The SOFFA data acquisition method differs from typical Fourier Block-Filtering by two important factors: (i) Each segment is a new collection of data with field- or frequency-stepped correlated signal and uncorrelated noise and (ii) each segment overlaps the next segments for a total of *m* overlapping segments. Due to oversampling, the spectral bandwidth of the spectrum in each segment is reduced, see Fig. A1 comparing spectra *ii* and *iii*.

Fourier block-filtering is illustrated in Fig. 2. Each segment is convoluted with the filter H[*ω*], such that

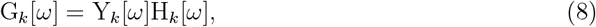

where Y_*k*_[*ω*] is the Fourier transform (Fourier space *ω*) of the discrete signal and the convolution is G_*k*_[*ω*]. In this work, a fixed FIR Gaussian filter is used, see Eq. 3. However, it should be noted that these filters do not have to be fixed and can vary at each *k* value, or be more sophisticated, such as adaptive averaging techniques.[25, 26]

Spectral leakage occurs when the chosen Gaussian standard deviation *σ*_*n*_ is more narrow than the spectral bandwidth of the desired signal. As the length of *L* is increased for each segment, the OSR must be increased to maintain the same spectral bandwidth. However, in practice this is impractical and, instead, the standard deviation of the filter must be modified to ensure no spectral leakage. By choosing to adjust *σ*_*n*_, the spectral bandwidth will approach that of the whole spectrum as *L* approaches the total length of 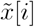 (*i* = 0, 1, …, *N*) and the signal-to-noise improvement will approach that of filtering after concatenation (Post-Add Filter) as employed by Hyde *et. al*.[4]

Since the optimum windowing method for the SOFFA algorithm is rectangular, abrupt discontinuities may exist in field space as the data is collected segmentally. Discontinuities cause Sinc-like features in Fourier space. However, Fourier filtering each segment with an FIR Gaussian filter removes these discontinuities and produces smooth overlap and concatenation. Filtering each segment separately removes the need for filtering the periodic noise associated with the step as in Ref. [4], which is required if the spectrum is filtered after concatenation as in Post-Add Filter.

As *m* increases the length of the tail further overlaps with the head of multiple segments resulting in averaging.

### 3.3 Segment-Overlap Averaging

Once each segment is successfully filtered, the segments must be concatenated together. Due to the use of a rectangular window, the segments can be concatenated in Fourier space, to save processing time, and a single inverse Fourier transform (IFFT) is needed to obtain the discretized spectrum. Because the data is zero filled and aligned, this is done simply by summation

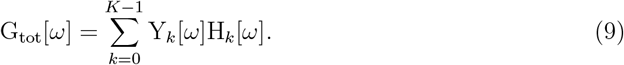

Since each of the segments has a portion of the signal and non-coherent noise that lies within the spectral bandwidth (low frequency), an additional signal-to-noise improvement of 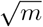 is expected by averaging the overlapped segments.

Finally, the first and last *L* points are called the head and tail, respectively. Since the head and tail do not have a complete segment-overlap dataset, they must be removed. As *m* increases, the head and tail regions may encroach on the magnetic resonance signal. In practice, for large values of *m* the total sweep width should be increased to accommodate for the removal of the head and tail region.

### 3.4 Decimation

The total number of points in the final concatenated spectrum *y*_tot_[*i*] is a function of the oversampling rate, the step size, and the number of steps. In order to generate a SOFFA acquired spectrum with a conventional number of points, decimation is used. Decimation acts as a moving average while reducing the total number of points. Herein a nearest neighbor windowing approach is used where *M* number of points are averaged together and replaced by one new point, where *M* is the re-sampling ratio.

### 3.5 Summary

The aforementioned processes provide a framework for the SOFFA data acquisition method. It should be noted that if the filter H[*ω*] is constant, as defined in this work, the output *y*[*i*] will be equivalent if block-filtering is used or the oversampled data is filtered after concatenation, but before decimation. By filtering after decimation (Post-Add Filter), the filtering effectiveness is reduced compared to the SOFFA method. If the filter varies for each segment H_*k*_[*ω*] it is no longer possible to obtain an equivalent filtering after concatenation, but before decimation. [23, 27] Thus, the block-filtering approach employed by the SOFFA method provides more flexibility in filter design. Effectively, the SOFFA data acquisition method splits data processing into two categories: (i) high-frequency filtering and noise shaping and (ii) low-frequency averaging.

## 4. Results and Discussion

### 4.1 Simulated Gaussian Absorption Spectrum

Segmented data is simulated with Fourier filtering (*σ*_*n*_ = 30) performed after decimation (Post-Add Filter; •), and an increase in signal-to-noise is shown related to the length of the overlap *m*, shown in Fig. 4. The Post-Add Filter is similar to the segmented-NARS methodology which has been previously reported and analyzed. [4] In this study, Post-Add Filtered data does not perform better than traditional averaging. However, this simulated study only characterizes the signal-to-noise assuming all experiments take the same amount of time. Further practical improvements are expected when accounting for the step size, oversampling rate, scan time, and conventional averaging time. For example, it is possible to create experimental parameters where 50 steps have an overlap of *m* = 200 and take the same total measurement time of conventional averaging of 50 total averages.

**Figure 4:**
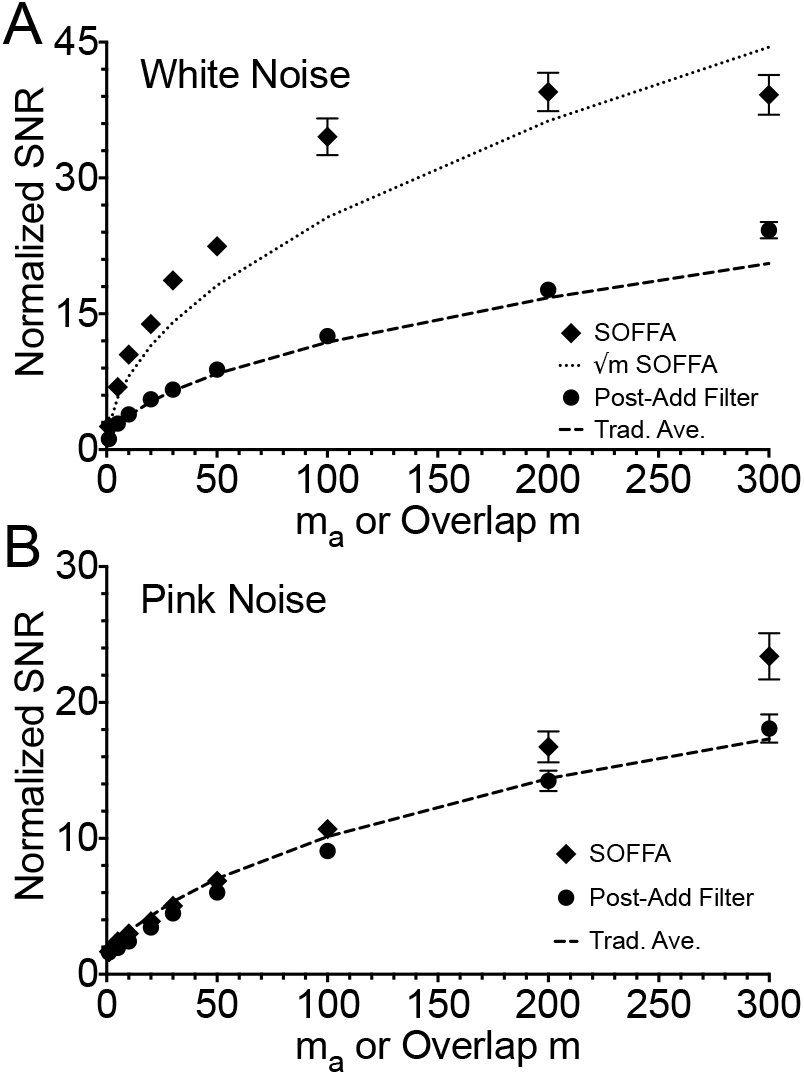
Signal-to-noise mean and 95% confidence interval (CI) of 100 simulations of a simulated Gaussian absorption spectrum with additive (A) white Noise and (B) pink Noise. Post-Add filter, with a fixed variance = 30 is shown as ^•^, while the SOFFA data acquisition method with the a variable variance is plotted as♦. Dotted lines are the expected 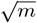 from averaging starting at SOFFA with no overlap *m* is one; 71.5. Dashed lines are the traditional averaging, which follows a strict 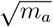 starting at a single scan *m*_*a*_ is one; 33.0. Data are normalized to the mean of the *m*_*a*_ = 1 value of the traditional averaging with white and pink noise, respectively. Data is tabulated in Table 1.

Finally, segmented data is simulated and processed with the SOFFA method. When *m* = 1, no overlap is present and the SOFFA method produces a factor of 2.2 (71.5:33.0) compared to filtering the conventional averaging (Trad. Ave.) spectra for white noise and a gain of 1.7 (15.2:9.1) for pink Noise. As the number of overlap segments increases the gain at *m* = 200 exhibits a factor of 2.4 (1101:467.3) for white noise or a potential of 1.4 (213.0:157.7) is realized when pink noise dominates. The improvement shown between the SOFFA *m* = 1 data and conventional averaging *m*_*a*_ = 1 is due to filtering oversampled segments and decimation of the concatenated whole spectrum. Data is tabulated in Table 1. It is clear from Fig. 4 that the SOFFA method (♦) performs exceedingly well on white noise compared to both conventional averaging (dashed) and the Post-Add Filter methods (•) of Ref. [4]. In Fig. 4A, a 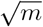 line is plotted normalized to *m* = 1 of the SOFFA method (dotted). As the overlap is increased and each dataset uses an optimum *σ*_*n*_, the improvement compared to 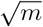 has a maximum at *m* = 200. This suggests there is an optimum overlap and filter combination that is a function of the spectral bandwidth. Additionally, as *m* increases the spectral bandwidth approaches that of the whole spectrum. When this occurs, filtering after decimation or with block-filtering becomes equivalent if H[*ω*] is constant.

When dealing with noise dominated by frequency dependent pink noise, the segmented methodology, in general, performs well. Pink noise is considered very difficult to filter since the spectral bandwidth is similar to the EPR spectrum. However, by effectively splitting the high frequency noise floor and averaging the low frequency, the signal-to-noise can be improved compared to traditional averaging and filtering, shown in Figs. 4B. Further improvement reducing pink noise is feasible by including adaptive averaging techniques as the filter H_*k*_[*ω*]. [25, 28, 26]

### 4.2 Continuous-Wave EPR

A series of CW EPR spectra of *apo* [FeFe]-hydrogenase from *Chlamydomonas reinhardtii* of varying concentrations were collected using a bismuth germanate (Bi4(GeO4)3, BGO) dielectric resonator retrofitted in a Bruker MD5 housing. [29] The *apo* protein has one reduced [4Fe-4S]^+^ cluster with an effective S=1/2. Three concentrations of 1 mM, 100 *µ*M, and 10 *µ*M were prepared and confirmed with UV-VIS by measuring the characteristic 400 nm feature indicating relative amounts of [4Fe-4S]. [18] The conventional-CW spectra were collected by a 4 minute scan of 4096 points over 100 mT. The 1 mM concentration spectrum, shown in Fig. 5A spectrum *i* is intended to be used as a reference. Both the 100 *µ*M and 10 *µ*M concentrations were collected with 25 averages for a total time of 100 minutes, shown in Fig. 5A spectrum *ii* and *iii*, respectively. Both CW spectra were background subtracted to remove background from the shield. All CW data was filtered with a *σ*_*n*_ = 75. The 100 *µ*M CW spectrum yielded a signal-to-noise of 181.8, while the 10 *µ*M CW spectrum was calculated at 16.5.

**Figure 5:**
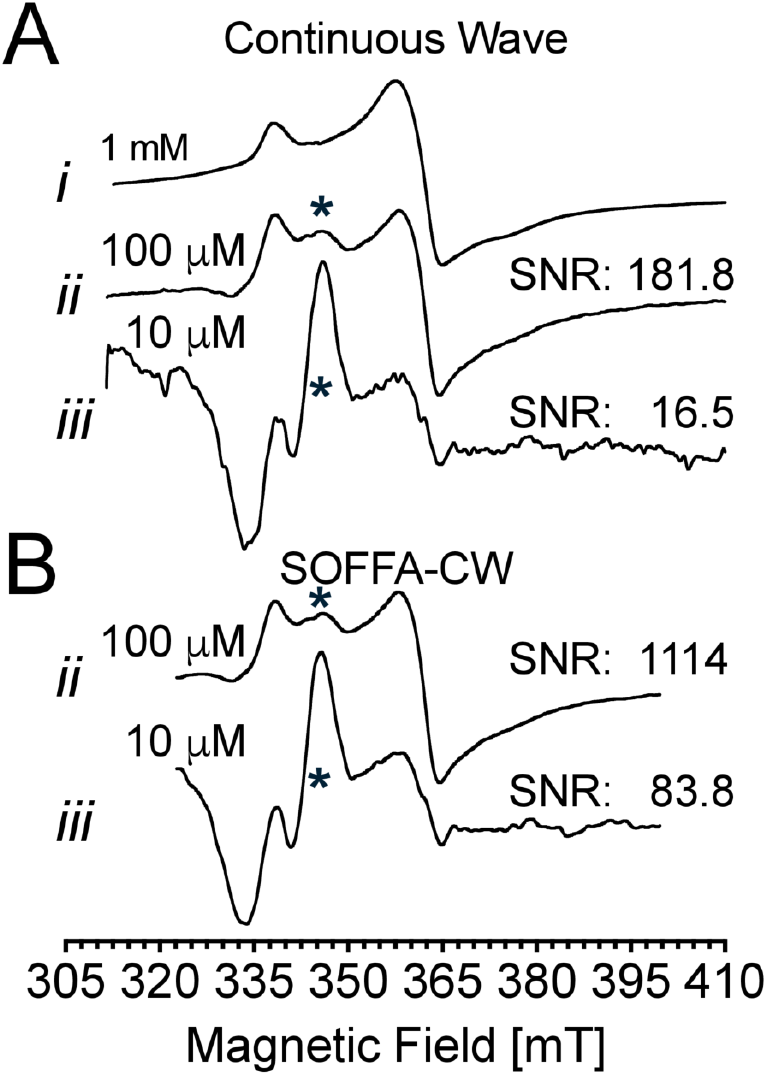
CW spectra were collected on an Bruker E5180 X-band bridge with a dielectric resonator (9.70 GHz) of the reduced [4Fe-4S]^+^ of *apo* [FeFe]-hydrogenase from *Chlamydomonas reinhardtii* at 1 mM, 100 *µ*M, and 10 *µ*M concentrations, spectrum *i, ii*, and *iii*, respectively. The reference spectrum of 1 mM concentration was collected over 100 mT, 4 minute scans averaged 9 times with 4096 points. The traditional CW experiment with 100 *µ*M concentration was collected over 100 mT, 4 minute scans averaged 25 times with 4096 points for a total time of 100 minutes. Both CW spectra were filtered with a *σ*_*n*_ = 75. The SOFFA-CW experiment with 100 *µ*M concentration was collected in 0.5 mT steps of 25 mT over 30 s and 4096 points for a total time of 100 minutes. The SOFFA-CW experiment with 10 *µ*M concentration was collected in 0.5 mT steps of 25 mT over 30 s and 4096 points for a total time of 100 minutes. For both SOFFA-CW experiments, a total of 200 steps were collected over a 100 mT total sweep. All segments were block filtered with a *σ* = 50 and with an *m* = 48. All spectra were collected at a temperature of 15 K and at an incident power of 0.63 mW with a field modulation amplitude of 0.5 mT.

A SOFFA-CW EPR spectrum was obtained by stepping the field 0.5 mT over 100 mT resulting in 200 steps of 25 mT sweep width (4096 points each). Each segment was collected over 30 seconds for a total time of 100 minutes. A signal-to-noise improvement of 6.1 (1114:181.8) is exhibited between the 100 *µ*M CW experiment and the SOFFA-CW experiment. The SOFFA-CW experiment with 10 *µ*M concentration was also collected in 0.5 mT steps of 25 mT sweep width over 30 s and 4096 points for a total time of 100 minutes. A total of 200 steps were collected over a 100 mT total sweep. The signal-to-noise ratio was calculated to be 83.8 and a signal-to-noise improvement of 5.1 (83.8:16.5) is exhibited for an average of 5.6 increase in concentration sensitivity. All SOFFA-CW spectra were block filtered with a *σ* = 50 and with an *m* = 48. The EPR feature at 340 mT is a background caused by residual sodium dithionite which is used to reduce the iron sulfur clusters of the *apo* [FeFe]-hydrogenase.

### 4.3 Non-adiabatic and Adiabatic Rapid Scan

Conventional CW data is compared to SOFFA-NARS of a 150 *µ*M of a site-directed spin-labeled Hemoglobin sample in 82% glycerol (18°C) at X-band (9.81 GHz), shown in Fig. 6. Experimental parameters are stated in the figure caption. This sample is in the slow tumbling regime of approximately 15 *×* 10^−9^ seconds rotational correlation time *τ*_*corr*_. After 16 averages of 2 min scans a CW signal-to-noise of 9.9 is realized. Adding a Gaussian filter increases the signal-to-noise by a factor of 4.7 (46.9:9.9). By processing the segmented-NARS data with the SOFFA filtering method, the collected pure-absorption signal has a signal-to-noise of 3890 (data omitted). In order to display the spectrum in a more conventional form, the first derivative is computed using an MDIFF pseudo-modulation of 0.1 mT. A signal-to-noise increase of 10.3 (483.3:46.9 compared to the filtered CW data) was achieved for the same measurement time of 32 minutes. The zoomed inset is the first 500 points of the NARS signal multiplied by a factor of 50. Since NARS uses a balanced mixer and A/D converter, the dispersion signal was also collected for no increase in measurement time (data omitted). The NARS linear background was baseline corrected in Xepr.

**Figure 6:**
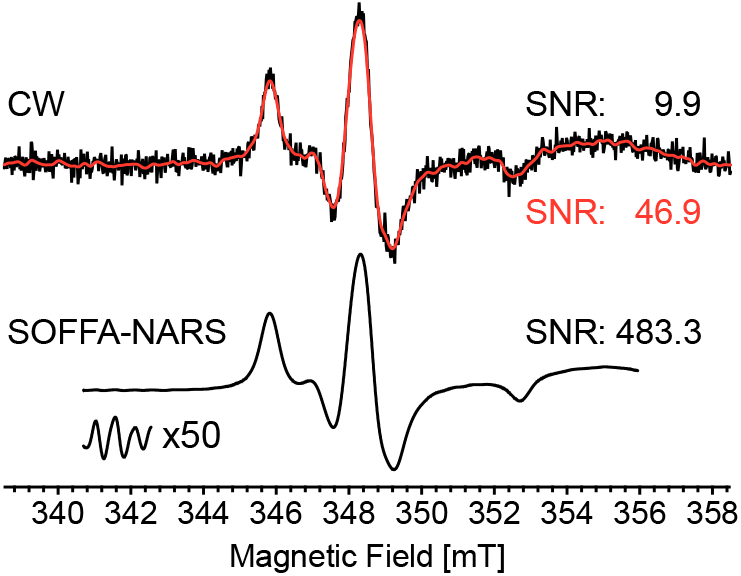
A 150 *µ*M site-directed spin-labeled Hemoglobin sample in 82% glycerol at 18C is collected with conventional-CW EPR in a five-loop–four-gap loop-gap resonator at X-band (9.81 GHz) using 0.1 mT field modulation at a rate of 100 kHz, 5 ms time constant, 16 averages over 20 mT (2 min scan) at 1 mW of microwave power yielding a signal-to-noise of 9.9. A Gaussian filter is applied to the CW data yielding a signal-to-noise of 46.9. SOFFA-NARS data is collected with 0.25 G steps (*s*) an overlap of *m* = 40 using a trapezoidal field ramp of 11.24 kHz with 6 *µ*s flat rate and an amplitude of 0.5 mT. Processing the data with the SOFFA method results in a signal-to-noise of the pure-absorption signal (not shown) of 3890. An MDIFF pseudo-modulation of 0.1 mT field modulation equivalent was used to display the spectrum in a more conventional form, yielding a signal-to-noise of 483.3. The zoomed inset is the first 500 points of the NARS signal. Data was collected on a custom-built X-band bridge dedicated to segmented-NARS spectroscopy. A NARS dispersion signal was also collected (not shown). Both experiments took approximately 32 minutes each.

As another example, an RS experiment was performed on a minuscule amount of LiPC at 2 mW, shown in Fig. 7. The CW data is shown with 16 averages over 0.1 mT (1 min scan) at 6.3 *µ*W with a modulation amplitude of 0.025 mT. A typical RS experiment of 1 mT sweep at a 100 kHz sinusoidal rate was performed and 2^16^ on-board-averages were averaged 324 times at 2 mW. A second spectrum was obtained by using a 1 mT sweep at a 100 kHz sinusoidal rate and 2^16^ on-board-averages were averaged 9 times at 2 mW. The field was stepped in 100 steps of 0.02 mT for a total sweep width of 2 mT and an overlap of *m* = 30.

**Figure 7:**
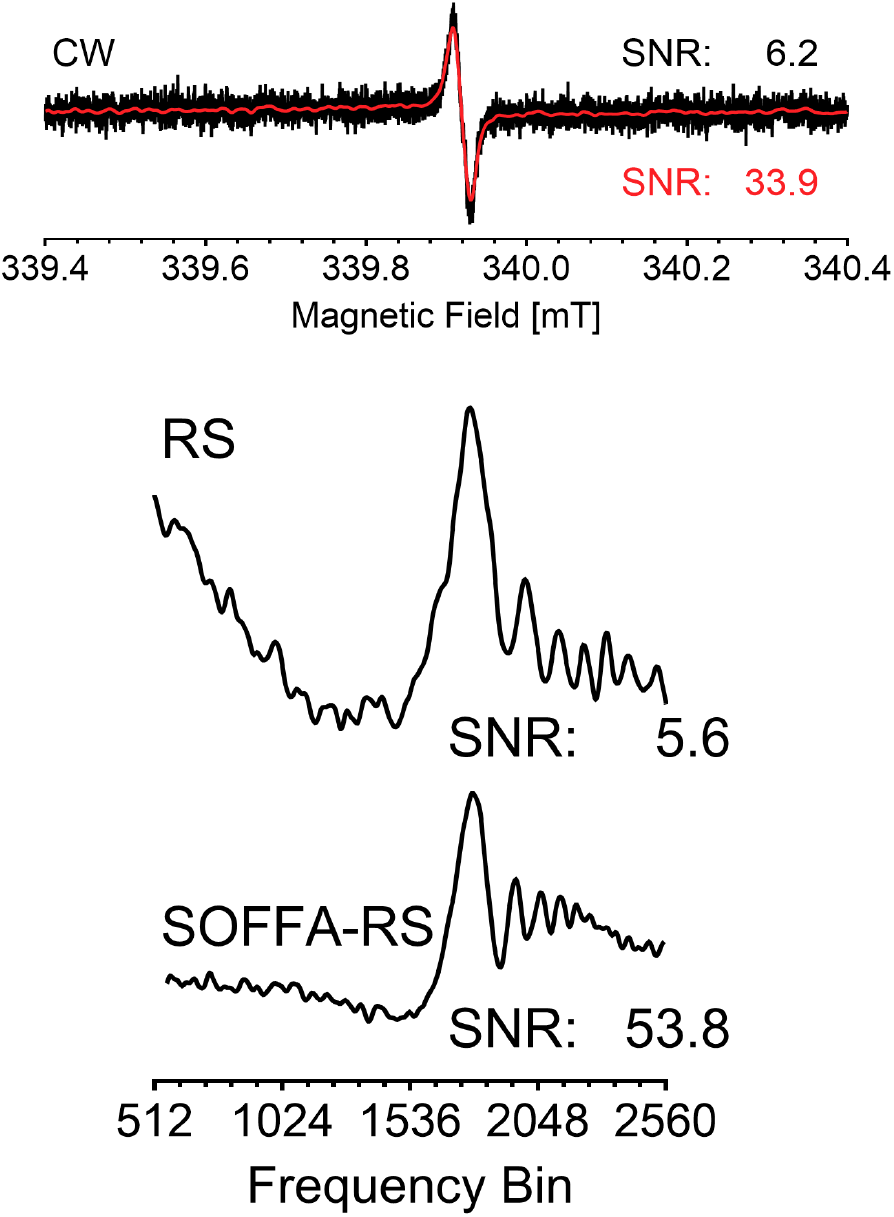
A lithium phthalocyanine (LiPC) is collected with conventional-CW EPR in a Bruker dielectric resonator at X-band (9.55 GHz) using 0.1 mT field modulation at rate of 100 kHz, 14 ms time constant (4096 pts), 16 averages over 1 mT (1 min scan) at 6.3 *µ*W of microwave power yielding a signal-to-noise of 6.2. A Gaussian filter is applied to the CW data yielding a signal-to-noise of 33.9. The RS signal was obtained by a 1 mT sweep at a 100 kHz sinusoidal rate and 2^16^ on-board-averages were averaged 324 times at 2 mW. A signal to noise of 5.6 is calculated. The SOFFA-RS spectra were obtained by using a 1 mT sweep at a 100 kHz sinusoidal rate and 2^1^6 on-board-averages were averaged 9 times at 2 mW. The data was stepped with 100 steps of 0.02 mT for a total sweep of 2 mT with *m* = 30. A signal-to-noise of 53.8 is calculated. No background was removed from both RS spectra to highlight the significantly decreased background in the SOFFA-RS spectrum. Traditional and segmented rapid scan experiments were performed using the methods described in Möser *et al*. [11] on a Bruker E580 X-band bridge. A RS dispersion signal was also collected (not shown). All experiments took approximately 16 minutes each.

All RS measurement times were approximately 16 minutes.

A factor of 9.6 (53.8:5.6) in signal-to-noise is achieved using segmented-RS and the SOFFA filtering method (SOFFA-RS) compared to traditional RS. The signal improvement is not only demonstrated by the reduction of noise, but also the increase in the modulation depth of the FID-like response. Background subtraction was not used for either dataset in order to highlight the advantage of the SOFFA method. Such a background in traditional RS could be removed by spectral fitting, deconvolution [7], specially-designed resonator probes [30], or background subtraction. [6] However, the use of background subtraction would increase the signal acquisition time by a factor of 2 and the noise displayed between pts 2304 to 3200 would most likely not be corrected. Whereas the SOFFA-RS dataset can be background subtracted from the first collected segment which is typically off-resonance and other noise is averaged during concatenation of the overlapping segments. As demonstrated with the SOFFA-RS, the inherent background-subtraction combined with concatenation of many segmented steps significantly reduces low-frequency background signals for the same measurement time.

Note that this experiment is not optimized to highlight the advantage of RS spectroscopy. Typically, one would further increase the incident power compared to the CW spectrum to arrive at the same ∂*B*_1_*/*∂*t* at resonance. After processing, the segmented scan data per the methods of Tseytlin *et al*.[16] or Joshi *et al*.[6] the slow-scan pure-absorption CW spectrum can be obtained.

### 4.4 Practical Considerations

The signal-to-noise ratio of EPR is fundamental to the usefulness of the technique. Improvements in EPR fall into two categories: technical developments, such as resonator and spectrometer advances, multiharmonic data collection[31] and advanced averaging and filtering techniques, such as, wavelet analysis[32] and adaptive averaging[25, 26] techniques. One must take care that any new method does not over-filter and deliver a distorted EPR spectrum.

In addition, the evaluation of new technology must be done with both random Gaussian noise and correlated 1/f noise under consideration. As describe earlier, Gaussian noise is characterized by the uniform power spectral density in the whole frequency space, plotted in Fig. A1B(vi). Correlated noise (“Pink” noise), as shown in Figure A1(v), where a significant portion of the power spectral density is located in low frequencies (1/f) that overlap with the power spectral density of the EPR signal. It is important to note that “Pink” noise power spectral density is a function of scan time and can be thought of as a measurement of system stability. [1] Because of this, changing the sweep speed changes the correlated noise of the system.

In this experiment, the ^13^C lines of 10 *µ*M TEMPOL have been highlighted by using traditional averaging, shown in black, by scanning 400 times over a 273 minute time period. In contrast, the same signal-to-noise ratio can be achieved in only 20 minutes (red) without loss of the subtle ^13^C lines.

Since the power spectral density of 1/f noise is similar to the EPR signal, both correlated noise and EPR signal have similar shape in the field domain making fitting subtle spectral features difficult without significantly improving the signal-to-noise ratio of the EPR signal with hardware improvements since excessively long averaging will not average out 1/f noise. As an example, the EPR spectra of Fig. 8A was corrupted with 1/f noise by averaging randomly generated correlated noise and adding it to the EPR spectrum. The noise was implemented in the frequency domain following a 1/f power law, establishing a crucial framework for realistic noise modeling in EPR experiments through a systematic process of creating frequency spectra, scaling amplitudes inversely with frequency, and adding random phase components. The implementation maintains mathematical rigor by ensuring conjugate symmetry in the frequency domain, a necessary condition for reconstructing physically meaningful real signals when transforming back to the field domain. The resulting spectrum for continuous wave is shown in black in Fig. 8B.

**Figure 8:**
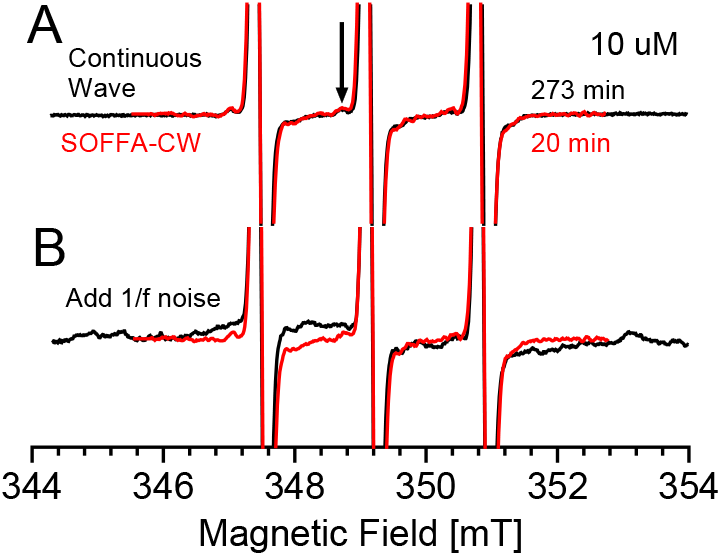
A 10 *µ*M TEMPOL spectrum is recorded using A) traditional CW averaging for 273 minutes (400 averages; black) and using SOFFA-CW in 20 minutes (red). On ^13^C feature is highlighted. The continuous wave experiment was collected with 10 mT sweep of 1024 points over 41.94 seconds with 400 averages, 20.48 ms time constant, at 3.2 mW, 0.1 mT field modulation at 100 kHz. SOFFA-CW was performed with 3 mT sweep of 8192 points over 10.39 seconds, with 100 steps of 0.1 mT each, 1.28 ms time constant, at 3.2 mW, 0.1 mT field modulation at 100 kHz. B) Phase noise was averaged for 400 averages and added to the continuous wave data (black) and phase noise was added onto each of the 100 steps of the SOFFA-CW (red).

For SOFFA, the randomly generated correlated noise was added to each of the 100 segments and averaging was performed through the SOFFA algorithm. Since the continuous wave EPR experiment and the SOFFA-CW experiment have different sweep times (41.49 s and 10.39 s, respectively) the frequency content of the correlated noise is different and it taken into account in this experiment. The resulting spectrum for SOFFA-CW is shown in red in Fig. 8B. From this, it is clear that the subtle ^13^C lines of the 10 *mu*M TEMPOL are lost in the continuous wave EPR spectrum and remain in the SOFFA-CW. The SOFFA method demonstrates enhanced EPR spectral features by not only enhancing the signal-to-noise ratio by a factor of 3.2 compared to traditional continuous wave measurements (1203:367), but also effectively mitigating phase noise and 1/f correlated noise, thereby preserving subtle spectral features such as ^13^C lines and enabling more reliable spectral fitting even in challenging low-concentration samples.

Finally, the version of Bruker Xepr (v.2.6b-160) used to collect the SOFFA spectra is not optimized for segmented data acquisition. Future work will implement segmented-NARS and RS using the Xepr Python API interface to decrease the latency associated with large datasets. A signal-to-noise improvement is expected by increasing the number of averages for the same total data-collection time. Despite current software limitations, the spectra collected in Figs. 5 and 7 were performed on a commercial Bruker E580 instrument with no hardware modifications. Further improvement in segmented-NARS and RS is expected by using trapezoidal sweeps and processing both the sweep-up and sweep-down data. [3, 33]

## 5. Applications

In this section, proposed applications are detailed where the SOFFA data collection method will be useful.

### 5.1 Segmented Continuous-Wave EPR

A segmented-CW EPR experiment processed by the SOFFA method (SOFFA-CW) is performed by collecting oversampled data from a standard lock-in phase-sensitive detector over a fraction of the spectrum and systematically stepping the center field to obtain the whole signal. The field modulation amplitude, time constant, and scan rate can all be adjusted to optimize the oversampled segments.

However, the spectral bandwidth of a CW experiment will always be larger compared to the pure-absorption spectral bandwidth due to higher frequency-content associated with the derivative-like shape from field modulation and phase-sensitive detect. When performing a segmented-CW experiment, using a large time-constant in a phase-sensitive detector acts as a low-pass filter, which may reduce the effectiveness of the SOFFA filtering. Collecting data with a low time-constant and applying the Fourier filter in post-processing provides more flexibility.

### 5.2 Field- or Frequency-Swept Segmented Non-adiabatic and adiabatic Rapid Scan

Field- or frequency-swept segmented-NARS and RS spectroscopy processed by the SOFFA method (SOFFA-NARS and SOFFA-RS) is performed by stepping the static magnetic field with a frequency/field sweep smaller than the total spectrum. The step size should be set so significant (*L >* 20*n*) overlapping occurs.

For NARS spectroscopy, pure absorption is directly collected. Pure absorption has the optimal spectral bandwidth for the SOFFA data acquisition method. In comparison, rapid scan has an increase of spectral bandwidth due to the FID-like frequency oscillations and may require over-coupling of the resonator to avoid filtering due to the resonator bandwidth. However, once RS data is collected, the pure absorption could be directly calculated by performing the deconvolution algorithm of Tseytlin *et al*. [16] block-wise before filtering and concatenation.

The implementation of multiple time averaging (MTA) is standard in NARS and RS spectroscopy due to the rapid collection of data at the rate of the field/frequency sweep.[14, 15] MTA is known to filter low frequency and shot noise associated with environmental and physiological noise. Since the data is collected rapidly, such noise is not readily added to the total signal.[14] However, as rates and amplitudes increase, the need to design resonators that exhibit low eddy-current forces (vibrational noise) is needed.

It should be noted that the first field-stepped RS spectra were obtained by Eaton and colleagues.[5] However, segmented-RS differs from the field-stepped RS spectra described therein. The work of Ref. [5] stepped the field while the whole RS spectrum was excited and then averaged together. In the current work, the field is stepped from an off-resonance position through the EPR signal and again to off-resonance.

No change in the collection of a NARS spectrum is needed to implement the SOFFA-NARS method.

### 5.3 Frequency-Stepped Segmented NMR and MRI

It has become common to use frequency stepping to obtain a broadband [34] or ultrawideband [35] NMR spectrum for exotic [36, 37, 38] and quadrupolar nuclei.[39] Such frequency-stepping techniques have also been extended for use with magic-angle spinning.[40] However, the use of frequency stepping could be configured for narrow-band acquisition in order to create a series of spectra that are segmented in frequency. Such a series of frequency-stepped segmented NMR spectra could be filtered by the SOFFA method and concatenated. Conceptually, this can be applied to two-dimensional experiments and experiments with gradients, such as MRI.

### 5.4 General Usage

Collection of SOFFA data requires the user to be able to step the magnetic field in known quantities (mT) and relate them to a set number of points. The parameters for a traditional CW experiment and SOFFA-CW experiment are summarized in Table 2. In modern EPR software (*e.g*., Bruker XEPR, Bruker MS5000, SpecMan) this is straight forward to calculate from the number of points that are set during configuration. In general, a new sample would require a traditional CW spectrum to be performed to know the full magnetic field width (Sweep Width) of the EPR signal.

**Table 2:** Comparison of parameters required for a traditional CW experiment and a SOFFA-CW experiment. The traditional time constant (RC filter) should be set as low as possible with SOFFA-CW. Total scan times do not take into account overhead (*e.g*., field flyback time, program lagging, etc.

|  | Traditional CW | SOFFA-CW |
| --- | --- | --- |
| Sweep Width ($t$) | 100 mT | 100 mT |
| Power | 0.6325 mW | 0.6325 mW |
| Mod. Amp. | 0.5 mT | 0.5 mT |
| No. of pts. ($N$) | 4096 | 4096 |
| Scan Time ($\tau$) | 240 s | 30 s |
| Aves. | 25 | 1 |
| Time Constant | 58.59 ms | - |
| Filter ($\sigma$) | - | 50 |
| Seg. Width ($g$) | - | 25 mT |
| Seg. Step ($s$) | - | 0.5 mT |
| Total Exp. Time | $t \times \text{Aves.}$<br>6000 s | $\tau \times (t/s)$<br>6000 s |

For a SOFFA-CW experiment, the user creates a new experimental setup that is separate from the traditional CW experiment. In the example of Fig. 5, another CW experiment (segment experiment) is chosen on Bruker XEPR and the user sets the spectrometer configuration to start at the lowest magnetic field. In this example, the SOFFA-CW segmented sweep width (Seg. Width) is chosen to be a quarter of the total typical CW sweep width (25 mT) and the number of points is set to be 4096. The time constant should be set to the lowest available, since we do not want to pre-filter the data for a SOFFA-CW experiment. If desired, the final SOFFA-CW signal can be filtered by an RC time constant for comparison to traditional CW.

In this example, the experimental scan time (30 s) of the segment experiment is set to 1*/*200th of the total time (including averaging) of the traditional EPR experiment to ensure the SOFFA-CW experimental time matches that of the total experimental time of the averaged traditional EPR spectrum. Further averaging of the series of segment experiments that make up a SOFFA-CW spectrum is always possible. However, the spectral overlap between segments means that there is already averaging built in (*m* of 48). Next, the ProDEL file is then loaded into the control and the field step (Seg. Step) is chosen in conjunction with the scan time to match 1*/*200th of the total field sweep (0.5 mT).

In the ProDEL code^1^, three variables need to be set. The first is the lowest magnetic field minus one step is placed in ‘value’, the final magnetic field stop is placed in ‘endParValue’, and the segment step is placed into the ‘parStep’ variable. Bruker XEPR then runs the ProDEL script on the active experiment.

Finally, the data is loaded into the Mathematica script^1^ and the appropriate variables are set. The filtering can be adjusted to ensure no broadening occurs (*σ* of 50). Future work will provide a python interface through the Bruker XEPR API to reduce latency and provide real-time filtering capabilities.

## 6. Conclusions

The SOFFA data acquisition method, introduced herein for segmented magnetic resonance spectroscopy, has been shown to decrease low-frequency background signals and improve the signal-to-noise ratio by a potential order of magnitude by effectively splitting high frequency filtering with low frequency averaging.

The use of the block-filtering approach employed by the SOFFA method provides a flexible framework for implementing complex filter designs. For example, frequency content could be analyzed and an appropriate filter could be chosen block-wise (*H*_*k*_(*ω*) where *k* = 0, 1, …, *K*; illustrated in Fig. 2) to maximize signal-to-noise while minimizing spectral broadening. Future work studying the effects of adaptive filtering and “spectral sensing” is underway in order to choose filter parameters without *a priori* information. [27, 41] Additionally, the SOFFA method lends itself to more advanced techniques, such as Wavelet analysis and filtering [32] or subband-multirate filtering [42], which may yield improved signal-to-noise and can be customized with *a priori* or real-time iterative calculation of the frequency content of the spectrum. Combining advanced filtering techniques with adaptive averaging [25, 28, 26] of the overlapping segments provides a powerful state-of-the-art digital signal processing toolkit for segmented spectroscopy. The SOFFA algorithm also lends itself to multi-harmonic detection.[43, 44]

Current studies using NARS and RS show spectral broadening due to edge effects of the applied sweeping field. In this work, the segment length *L* was kept small to remain in the more-linear portion of the sinusoidal sweep, thus negating this effect. However, this spectral broadening limits the size of *L*. This challenge can be addressed by using a trapezoidal sweep where the segments are collected during the linear rise or fall of the field/frequency sweep and the plateau is longer than the relaxation time of the spin system. This setup would minimize edge effects caused by abrupt changes of a triangular sweep or turning points of a sinusoidal sweep. Potentially, for RS, deconvolution of each segment block-wise before filtering is feasible. Future work will explore these possibilities.

As frequency-swept NARS and RS are more widely adopted, the SOFFA filtering technique introduced here may provide additional benefit by filtering and averaging signals from the frequency response of microwave components which create large and often quadratic backgrounds. [45, 46, 47] This has been demonstrated in Fig. 7, where the first segment contained no RS data and was used for a field-sweep background correction in SOFFA-RS. Such a background correction spectrum is taken separately for traditional RS and may produce much larger backgrounds due to the large eddy current constraints (or frequency response curves) seen in RS spectroscopy. SOFFA-RS may reduce the field sweep (frequency sweep) required to cover a spectrum.

Implementing SOFFA-CW by collecting segmented CW, an increase in concentration sensitivity of 6.9 is demonstrated for the same measurement time. In a SOFFA-NARS experiment, a factor of 10.3 signal-to-noise improvement, shown in Fig. 6, is achieved with no change in the experimental procedure described in the literature. [1, 4, 3] With RS, a segmented methodology must be adopted. However, once adopted, a factor of 9.6 signal-to-noise improvement is demonstrated in SOFFA-RS for the same measurement time, shown in Fig. 7. Further improvement compared to CW can be obtained in SOFFA-RS by increasing the incident microwave power to arrive at the same ∂*B*_1_*/*∂*t* at resonance. The SOFFA method is also useful for real-time segmented-NARS and segmented-RS processing, since the Fourier block-filtering can be easily parallelized on modern computers and displayed as the data is collected.

## 7. Acknowledgments

This work is funded by the European Union Horizon 2020 Marie Sk-lodowska-Curie Fellowship (Act-EPR; No. 745702), the Max Planck Society, and NIH NIGMS (no. R01GM149568-01). The author would like to thank Dr. Aaron W. Kittell and Prof. James S. Hyde for their initial work in the field of non-adiabatic rapid scan and Drs. Shannon A. Bonke and Joscha P. Nehrkorn for their helpful discussions. Dr. Constanze Sommer for the *apo* [FeFe]-hydrogenase sample. The author would also like to thank Profs. Wolfgang Lubitz and Dieter Suter for their continued mentoring and Drs. Edward J. Reijerse and Alexander Schnegg for their exceptional support, discussions, and encouragement.

## [Appendix A] Shape of Noise

Plots of simulated Gaussian spectra in field and Fourier space, Figs. A1A and A1B, respectively. A noiseless Gaussian is shown in Fig. A1 spectrum *i* and white noise is added to make the signal-to-noise approximately 4, shown in Fig. A1 spectrum *ii*. By filtering the noisy spectrum with a Gaussian filter (Fig. A1B; dashed-line) the noise outside of the spectral bandwidth (*f*_*cw*_) can be removed, shown in Fig. A1 spectrum *iv*. However, by oversampling, shown in Fig. A1 spectrum *iii*, the spectral bandwidth is reduced (*f*_*s*_) by the oversampling factor OSF and the noise power spectral density is reduced by the same amount. In order to mimic the methodology of oversampled segmented-data collection, the number of points remain the same, but the collected data is only a segment with a quarter of the field sweep. An example of pink noise is shown in Fig. A1 spectrum *v*. The power spectral density, shown in Fig. A1B illustrates the challenge with filtering pink noise: the concentration of power spectral density in the same region of the spectral bandwidth. Whereas white noise, shown in Fig. A1 spectrum *vi*, has a flat power spectral density response.

**[Figure A1:**
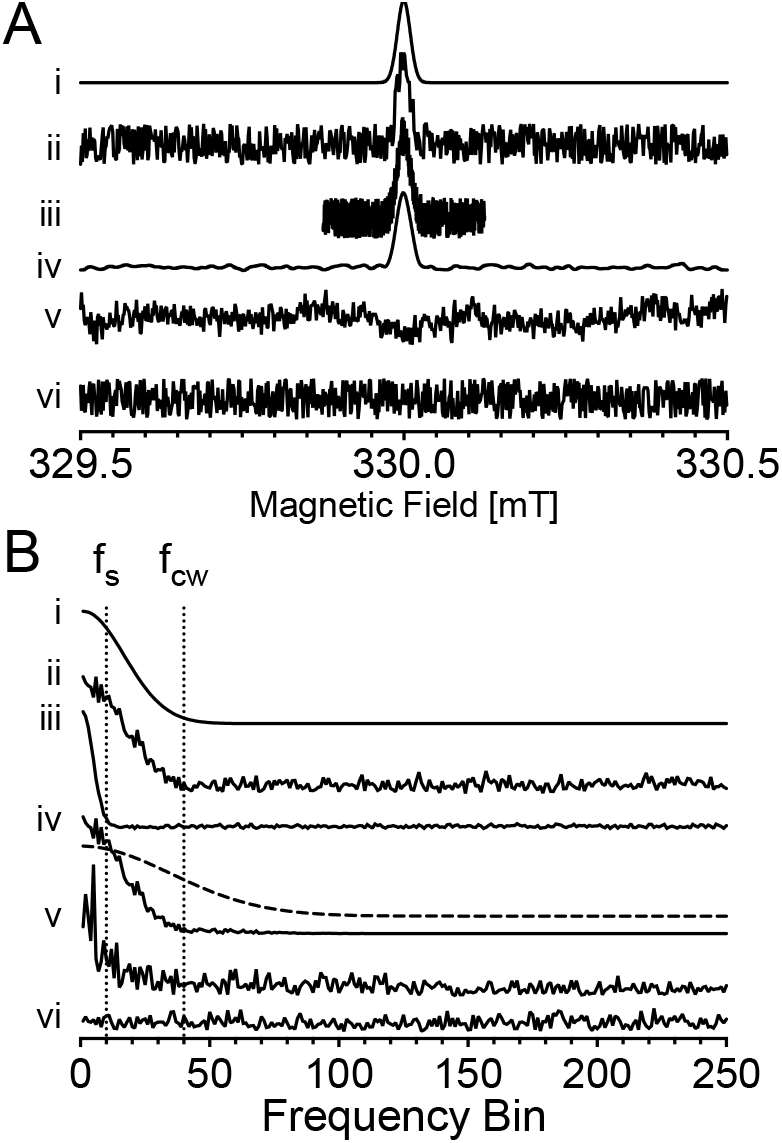
Simulated field and calculated corresponding Fourier transform employed to illustrate the details of spectral bandwidth. Simulated data are plotted in (A) Field space and (B) truncated Fourier Space to provide sufficient detail. Plotted is a (i) noiseless Gaussian spectrum, (ii) Gaussian spectrum with additive white noise, (iii) oversampled Gaussian spectrum with additive white noise (OSR = 4), (iv) Fourier-filtered Gaussian spectrum with Gaussian filter (dashed), (v) pink noise with noise floor, (vi) white noise.

## [Appendix B] Variable Definitions

The nomenclature of the Welch FFT procedure for power spectral density is adopted, due to the similarity of segmentation of a discretized signal. The variables used in this work are outlined in Fig. B1. [19]

**[Figure B1:**
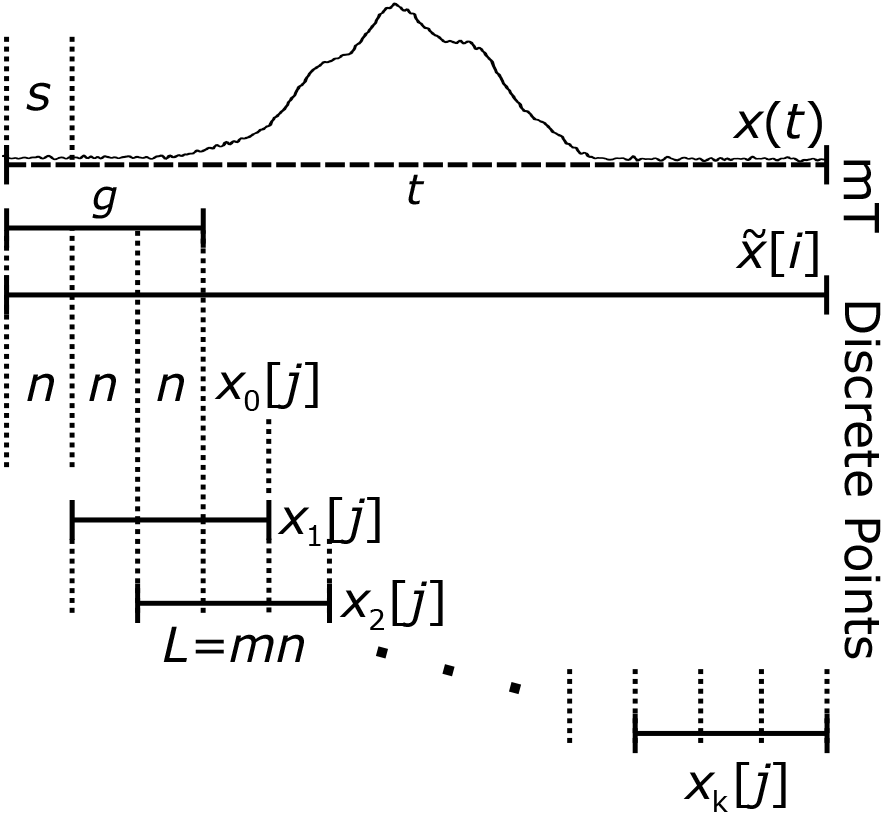
Illustration of spectral segmentation with parameters and variables.

## Footnotes

[1 The ProDEL program used to collect SOFFA-CW data can be found at the https://github.com/jsidabras/SOFFABruker.

